# A novel type of IRES having variable numbers of eIF4E-binding site and synergistic effect in RNA2 of WYMV isolates

**DOI:** 10.1101/2021.03.23.436552

**Authors:** Geng Guowei, Yu Chengming, Li Xiangdong, Shi Kerong, Yuan Xuefeng

## Abstract

Some viral proteins were translated in cap-independent manner via internal ribosome entry site (IRES), which ever maintained conservative characteristic among different isolates of same species of virus. However, IRES activity presented 7-fold of variance in RNA2 of wheat yellow mosaic virus (WYMV) HC and LYJN isolates. Based on RNA structure probing and mutagenesis assay, the loosened middle stem of H1 and hepta-nucleotide top loop of H2 in LYJN isolate synergistically ensured the higher IRES activity than that in HC isolate. In addition, the conserved top loop of H1 ensured basic IRES activity in HC and LYJN isolates. RNA2 5′-UTR specifically interacted with the wheat eIF4E, which was accomplished by the top loop of H1 in HC isolate or the top loop of H1 and H2 in LYJN isolate. Different IRES activity of WYMV RNA2 was regulated by different numbers of eIF4E-binding site and their synergistic effect, which was accomplished by the proximity of H1 and H2 due to the flexibility of middle stem in H1. It is represented a novel evolution pattern of IRES.

## Introduction

mRNA is generally translated via canonical ribosome scanning mechanism, in which 5′-cap of mRNA is responsible for firstly recruiting eIF4E followed by a series of translation initiation factors as well as 40S ribosome subunit to initiate translation (Marcotrigiano *et al,* 1997; Pestova *et al,* 2007; Sonenberg & Hinnebusch, 2009). However, some RNA viruses contained genomic RNAs without the 5′-cap and/or 3′-poly (A), which is essential for canonical ribosome scanning mechanism. Proteins of these RNA viruses were expressed through cap-independent translation manner, which can be mediated by two type of *cis*-elements including internal ribosome entry site (IRES) and/or cap-independent translation enhancer (CITE) (Kieft, 2008; Nicholson & White, 2011; Simon & Miller, 2013). IRES was located at 5′-upstream of corresponding open reading frame (ORF), while CITE was located at 3′-downstream of ORF. CITE was also termed as 3′-CITE, which could be delivered to 5′-upstream to regulate translation through long-distance RNA-RNA interaction (Simon & Miller, 2013) or interaction between ribosome 40S and 60S subunit in the case of TCV (Stupina *et al,* 2008; Stupina *et al,* 2011). Except to RNA viruses, host cellular mRNA also contained numerous IRES and 3′CITE (Baird *et al,* 2006; Ungureanu *et al,* 2006; Xue *et al,* 2015; Weingarten-Gabbay *et al,* 2016), which could be used to response specific and adverse condition.

IRES was firstly reported in picornavirus RNAs (Jang *et al,* 1988; Pelletier & Sonenberg, 1988). Although cellular mRNAs were also reported to contain IRES, detailed characteristic of IRES was mainly identified in RNA viruses especially animal RNA viruses. In general, IRESes in animal viruses had more complex structures than that of other resource (Kieft, 2008; Filbin *et al,* 2013; Hashem *et al,* 2013; Fern′andez *et al,* 2014). Based on the structure characteristic and mechanism promoting initiation involved in the various initiation factors and/or IRES *trans*-acting factors (ITAFs), IRESes from animal RNA viruses were classified into six types, including one type from the Dicistroviridae family and five types mainly from the members of the Picornaviridae family (Sweeney *et al,* 2012; Lozano & Mart′ınez-Salas, 2015). IRESes in plant viruses were mainly identified in members of the Potyviridae family such as Tobacco etch virus (TEV), Turnip mosaic virus (TuMV), Potato virus Y (PVY) and Wheat yellow mosaic virus (WYMV) (Zhang *et al,* 2015; Geng *et al,* 2020). The structure of IRESes in plant viruses was not complicated as that in animal viruses but usually had a weak secondary structure with few hairpins, which had unique characteristic from that of animal viruses (Zhang *et al,* 2015). It is implied the different evolution pathway for IRESes of different resource. Based on core structural features of IRESes, compounds or small molecules were designed to inhibit viral protein translation, which provided potential ways to control viral diseases (Seth *et al,* 2005; Gasparian *et al,* 2010; Dibrov *et al,* 2012; Boerneke *et al,* 2014; Lozano & Mart′ınez-Salas, 2015; Geng *et al,* 2020). Previous study on IRESes were focused on identification of structure characteristic, core *cis*-elements as well as *trans*-acting factors. Some RNA viruses had multi-segmented genome, in which each genome may have own IRES with special characteristic. In addition, IRES in different isolates of same virus may also have special characteristic. Study of the comparation and evolution among these IRESes should be helpful to identify the evolution of RNA viruses. However, variable IRES study among different isolates of same virus had few data.

Cap-independent translation of WYMV RNA1 has been regulated by dynamics equilibrium state of its IRES, which is mediated by discontinuous C-G base pairs (Geng *et al,* 2020). Meanwhile, this specific model of the IRES in WYMV RNA1 has been speculated to be conserved among different isolates due to the conserved 5′-UTR of different WYMV RNA1 (Geng *et al,* 2020). In contrast to high identity of the 5′-UTR in WYMV RNA1, the 5′-UTR of different WYMV RNA2 presented higher variance (Geng *et al,* 2017), which implied the potential difference in the cap-independent translation of different WYMV RNA2.In this study, variable IRES activity of 5′-UTR of different WYMV RNA2 were identified, which was mainly associated with different numbers of recruitment sites on wheat eIF4E and the synergistic effect between these recruitment sites.

## Results

### 5′-UTR of different WYMV RNA2 presented variable IRES activity with 7-fold difference, which can be enhanced by the corresponding 3′-UTR

To analyze and compare the effect of the 5′-UTR of WYMV RNA2 on translation, 5′-UTRs of three WYMV RNA2 isolates (HC with accession No. AF041041, TADWK with accession No. KX258950 and LYJN with accession No. KX258949) were respectively inserted upstream of the firefly luciferase (Fluc) ORF (Fig.1A and 1B), whose transcripts were performed *in vitro* translation using WGE system. Compared with the basic vector (F), 5′-UTR of three WYMV RNA2 isolates increased the translation level of Fluc to 10- to 70-folds in the absence of the 5′-cap, which presented high cap-independent translation activity (Fig.1B).

**Figure 1.**
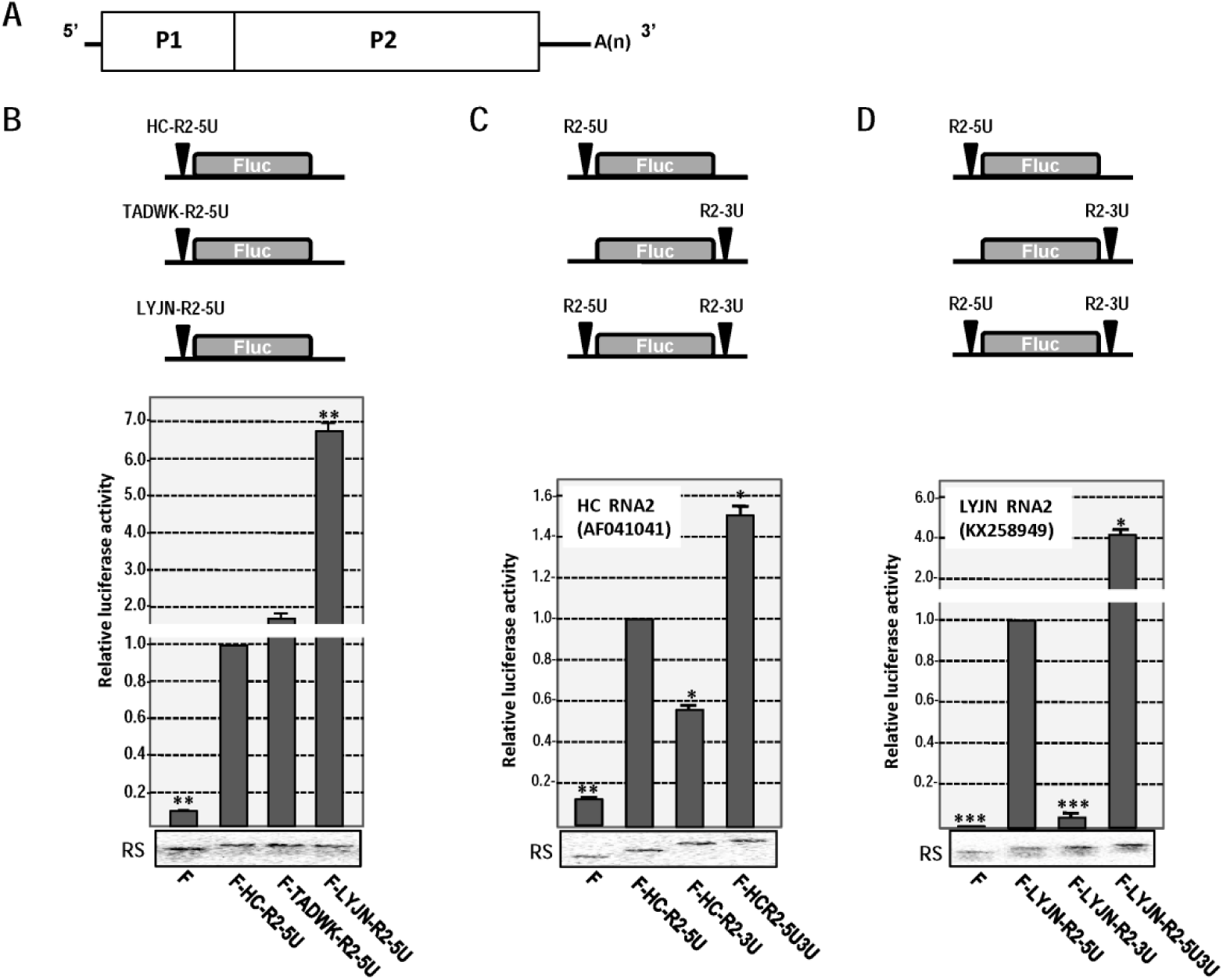
Effect of the UTRs in wheat yellow mosaic virus RNA2 on translation. **A. Genome organization of WYMV RNA2.** **B. Effect of the RNA2 5′-UTR of three WYMV isolates on translation**. Fluc, firefly luciferase; F, control vector; HC, Huangchuan isolate (RNA2 accession number: AF041041); TADWK, Taian isolate (RNA2 accession number: KX258950); LYJN, Linyi isolate (RNA2 accession number: KX258949); R2, RNA2; 5U, 5′-UTR. **C. Effect of the 5′-UTR and 3′-UTR of RNA2 of WYMV HC isolate on translation.** 3U, 3′-UTR. **D. Effect of the 5′-UTR and 3′-UTR of RNA2 of WYMV LYJN isolateon translation.** * indicates P <0.05, ** indicates P <0.01, *** indicates P <0.001; RS indicates RNA stability assay after 1.5 h *in vitro* translation.

Due to the remarkable 7-fold of difference of cap-independent translation activity, following assays were performed on RNA2 of HC isolate (Accession No. AF041041) and LYJN isolate (Accession No. KX258949). Presence of corresponding 3′-UTR enhanced the cap-independent translation activity of the 5′-UTR of WYMV RNA2 by 1.5-fold for HC isolate or 4.2-fold for LYJN isolate, which suggested the synergistic function of 5′-UTR and 3′-UTR on cap-independent translation (Fig.1C and 1D). When a long stem-loop (SL) was added upstream, the 5′-UTR of WYMV RNA2 still presented enhancement on cap-independent translation. For RNA2 of HC isolate, 5′-UTR (F-SL-HC-R2-5U) enhanced translation to 4.6-fold of that of control vector (F-SL) in the presence of upstream long SL (Fig.2A). For RNA2 of LYJN isolate, 5′-UTR (F-SL-LYJN-R2-5U) enhanced translation to 33-fold of that of control vector (F-SL) in the presence of upstream long SL (Fig.2B), It is suggested that 5′-UTR of WYMV RNA2 had IRES activity. In addition, 5′-UTR of WYMV RNA2 also enhanced cap-dependent translation by 4-5 folds, which was shown as the translation level of Cap-F-R2-5U vs Cap-F (Fig.2A and 2B). Taken together, RNA2 5′-UTR of HC and LYJN can not only enhance cap-dependent translation at similar level but also regulate cap-independent translation with 7-fold of difference.

**Figure 2.**
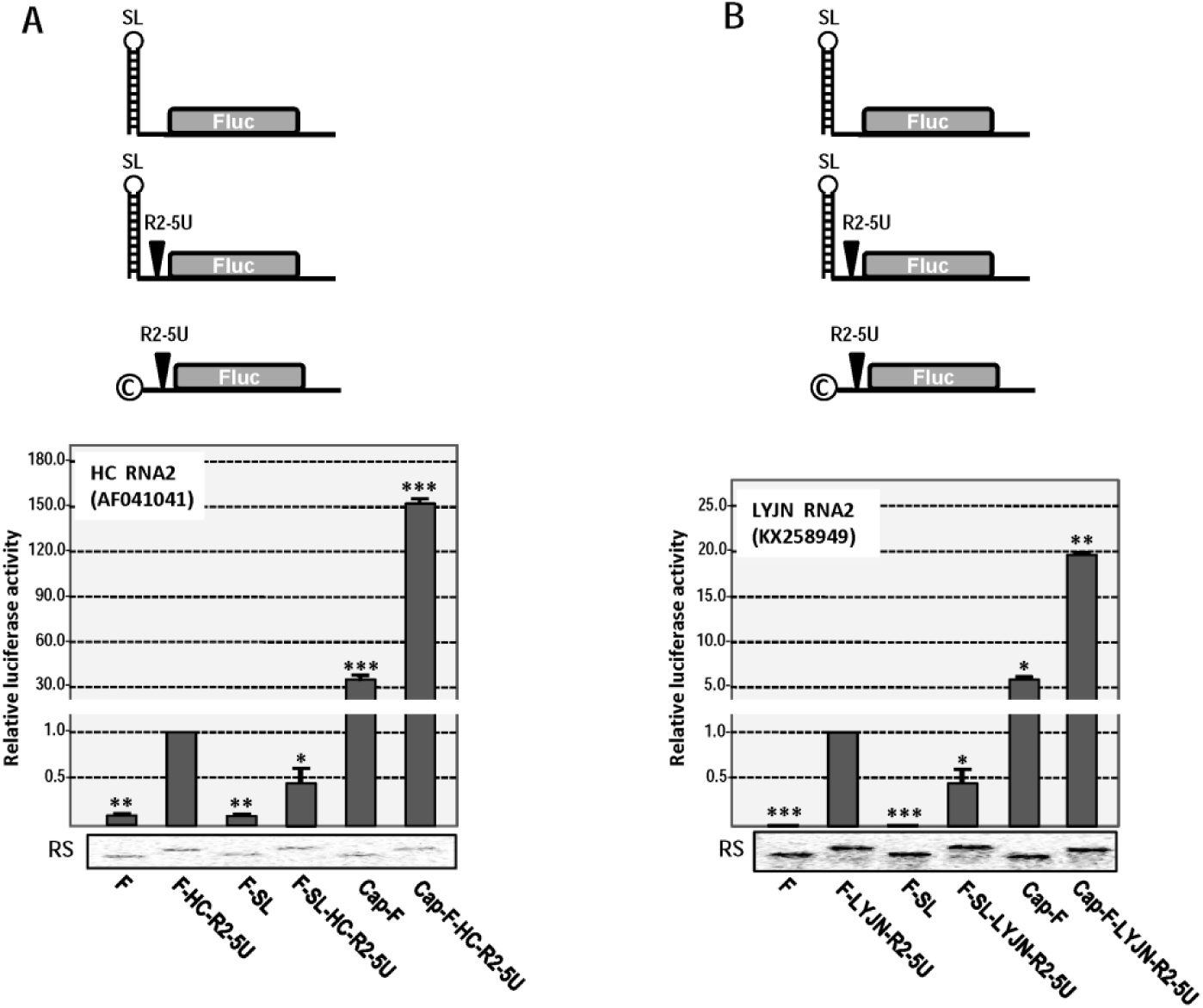
Effect of the RNA2 5′-UTR of WYMV HC and LYJN isolates on translation in the presence of additional upstream stem-loop or 5′-cap structure. **A. Effect of the RNA2 5′-UTR of WYMV HC isolate in the presence of additional upstream stem-loop or 5′-cap structure**. SL, stem-loop. **B. Effect of the RNA2 5′-UTR of WYMV LYJN isolate in the presence of additional upstream stem-loop or 5′-cap structure.** * indicates P <0.05, ** indicates P <0.01, *** indicates P <0.001; RS indicates RNA stability assay after 1.5 h *in vitro* translation.

To map the boundaries of IRES in the RNA2 5′-UTR of HC or LYJN isolate, a series of deletion mutants from the 5′-end or 3′-end with the 5′-UTR of RNA2 were constructed and performed *in vitro* translation (Fig.3A and 3B). After deletion of 20 nt from the 5′-end (F-HC-R2-5U-5′Δ20) or the 3′-end (F-HC-R2-5U-3′Δ20) in RNA2 5′-UTR of HC isolate, the translation of mutants remained about 85% or 78% of that of wt (F-HC-R2-5U) (Fig.3A). However, after deletion of 30 nt from the 5′-end (F-HC-R2-5U-5′Δ30) or the 3′-end (F-HC-R2-5U-3′Δ30) in RNA2 5′-UTR of HC isolate, the translation of mutants only have about 50% or 40% of that of wt (F-HC-R2-5U) (Fig.3A). Further 10 nt deletion from 5′-end or 3′-end caused a sharp loss of translation, it is suggested that the core region of IRES within the RNA2 5′-UTR of HC isolate was located between position 20 to position 151. Based on similar analysis, the core region of IRES within the RNA2 5′-UTR of LYJN isolate was located between position 30 to position 152 (Fig.3B).

**Figure 3.**
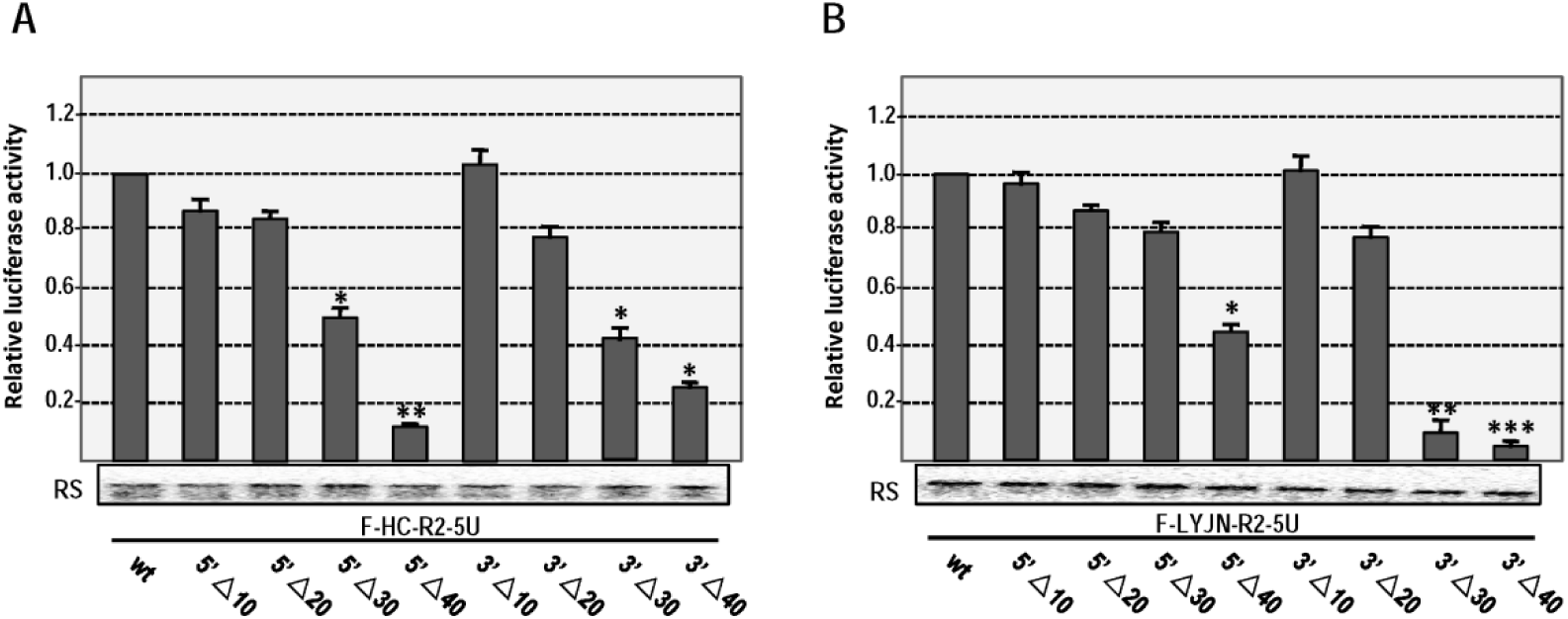
Effect of the deletion mutations in the RNA2 5′-UTRs of WYMV HC and LYJN isolates on translation. **A. Effect of the deletion mutations in the RNA2 5′-UTRs of WYMV HC isolate on translation.** **B. Effect of the deletion mutations in the RNA2 5′-UTRs of WYMV LYJN isolate on translation.** 5′Δ10, 5′Δ20, 5′Δ30, and 5′Δ40, respectively indicate the deletion of the terminal 10 nt, 20 nt, 30 nt and 40 nt from 5′-end of the RNA2 5′-UTR; 3′Δ10, 3′Δ20, 3′Δ30, and 3′Δ40, respectively indicate the deletion of the terminal 10 nt, 20 nt, 30 nt and 40 nt from 3′-end of the RNA2 5′-UTR. * indicates P <0.05, ** indicates P <0.01, *** indicates P <0.001; RS indicates RNA stability assay after 1.5 h *in vitro* translation.

### Different IRES activity of WYMV RNA2 5′-UTR was mainly associated with base-paring status of middle stem in hairpin H1 and nucleotide specificity of top-loop in hairpin H2

To identify the mechanism associated with the variable IRES activity of RNA2 5′-UTRs, the structures of RNA2 5′-UTR of HC and LYJN isolates were analyzed through in-line probing (Fig.4). Based on in-line cleavage pattern (Fig.4A and 4B), 5′-UTR of WYMV RNA2 had three hairpins (H1, H2 and H3), of which H1 and H2 were mainly located within the core region of IRES (Fig.3, Fig.4C and 4D). Based on the structure characteristic of RNA2 5′-UTR of HC and LYJN isolate, there are remarkable difference in two regions, which included base-pairing pattern of middle stem in H1 and nucleotide specificity of top loop in H2 (Fig.4C and 4D). In RNA2 5′-UTR of HC isolate (HC-R2-5U), middle stem in H1 presented firm base pairing between ^31^AAAGCUAU and ^75^AUAGCUUU (Fig.4A and 4C). In RNA2 5′-UTR of LYJN isolate (LYJN-R2-5U), middle stem in H1 failed to form base pairing between ^32^AAAACUAC and ^76^ACAGCUUU, which also loosened the base short stem in H1 (Fig.4B ad 4D). For top loop of H2, there was CCC in HC isolate and CCCUAUC in LYJN (Fig.4C and 4D). Other regions in H1 and H2 remained similar status and characteristic between HC and LYJN isolates. The difference in middle stem of H1 and top loop of H2 may be associated with the IRES activity between HC and LYJN isolates.

**Figure 4.**
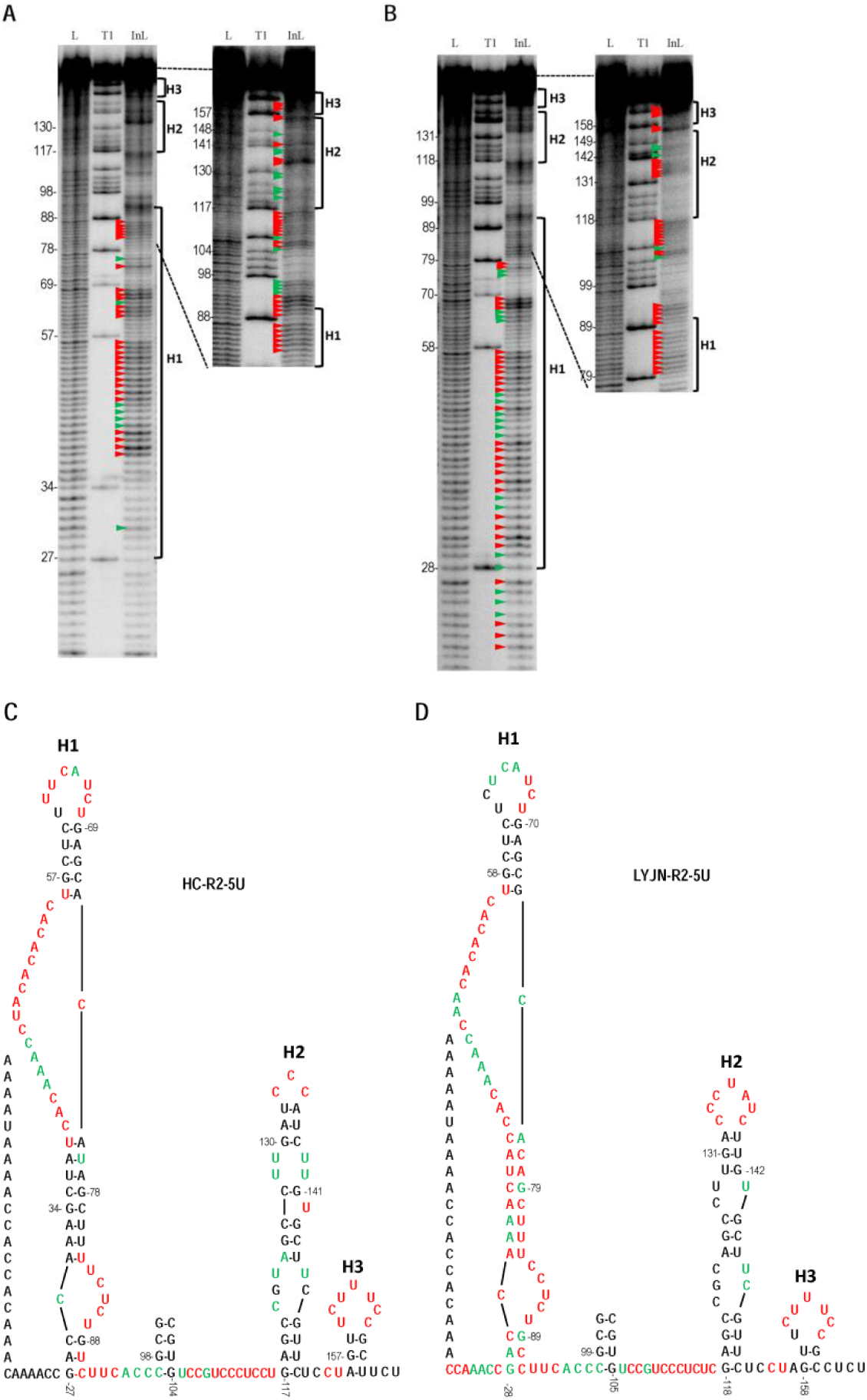
Structure probing of the RNA2 5′-UTR of WYMV HC and LYJN isolates. A. **In-line probing of RNA2 5′-UTR of WYMV HC isolate.** L: ladder generated from treatment with NaOH; T1: ladder generated from fragment denaturation and treatment with RNase T1, which cleaves at guanylates. InL: in-line cleavage products. Base numbering is at the left of the panel and positions of hairpins (H) are noted to the right. Red and green solid triangles point to strongly marked and weakly marked cleavage sites in the RNA2 5′-UTR. H1: hairpin 1; H2: hairpin 2; H3: hairpin 3. B. **In-line probing of RNA2 5′-UTR of WYMV LYJN isolate.** C. **RNA structure of RNA2 5′-UTR of WYMV HC isolate**. Black, green and red nucleotides respectively indicate none or inapparent, weakly marked and strongly marked cleavage based on the in-line probing pattern. Green and red nucleotides indicate the single-strand characteristics. D. **RNA structure of RNA2 5′-UTR of WYMV LYJN isolate.**

To identify the relationship between these two regions in H1 and H2 and remarkable IRES difference, mutual-mimesis types of mutants were constructed on RNA2 5′-UTR of HC and LYJN isolates (Fig.5). For RNA2 5′-UTR of HC isolate, mutant MS1 caused loosened status of middle and base stem in H1 and mutant ML1 changed the nucleotide of top loop in H2 as that of LYJN (Fig.5A). IRES activity of MS1 and ML1 of HC had respectively 2-fold and 3.2-fold of that of wt (F-HC-R2-5U) (Fig.5C). In addition, IRES activity of dual-mutant MS1+ML1 had 4.6-fold of that of wt HC (F-HC-R2-5U) and 68% of that of wt LYJN (F-LYJN-R2-5U) (Fig.5C). For RNA2 5′-UTR of LYJN isolate, mutant MS1 was designed to form firm base pairing of middle and base stem and mutant ML1 changed the nucleotide of top loop in H2 as that of HC isolate (Fig.5B). IRES activity of MS1 and ML1 of LYJN had respectively 50% and 37% of that of wt LYJN (F-LYJN-R2-5U) (Fig.5C). In addition, IRES activity of dual-mutant MS1+ML1 had 31% of that of wt LYJN (F-LYJN-R2-5U) and 2.1-fold of that of wt HC (F-HC-R2-5U) (Fig.5C). Taken together, it is suggested that base-paring status of middle stem in H1 and nucleotide specificity of top loop in H2 are two critical factors regulating the variable IRES activity of RNA2 5′-UTR. Loosened middle stem in H1 and hepta-nucleotide specificity in top loop of H2 were synergistically correlated with the higher IRES activity of RNA2 5′-UTR of LYJN isolate than HC isolate.

**Figure 5.**
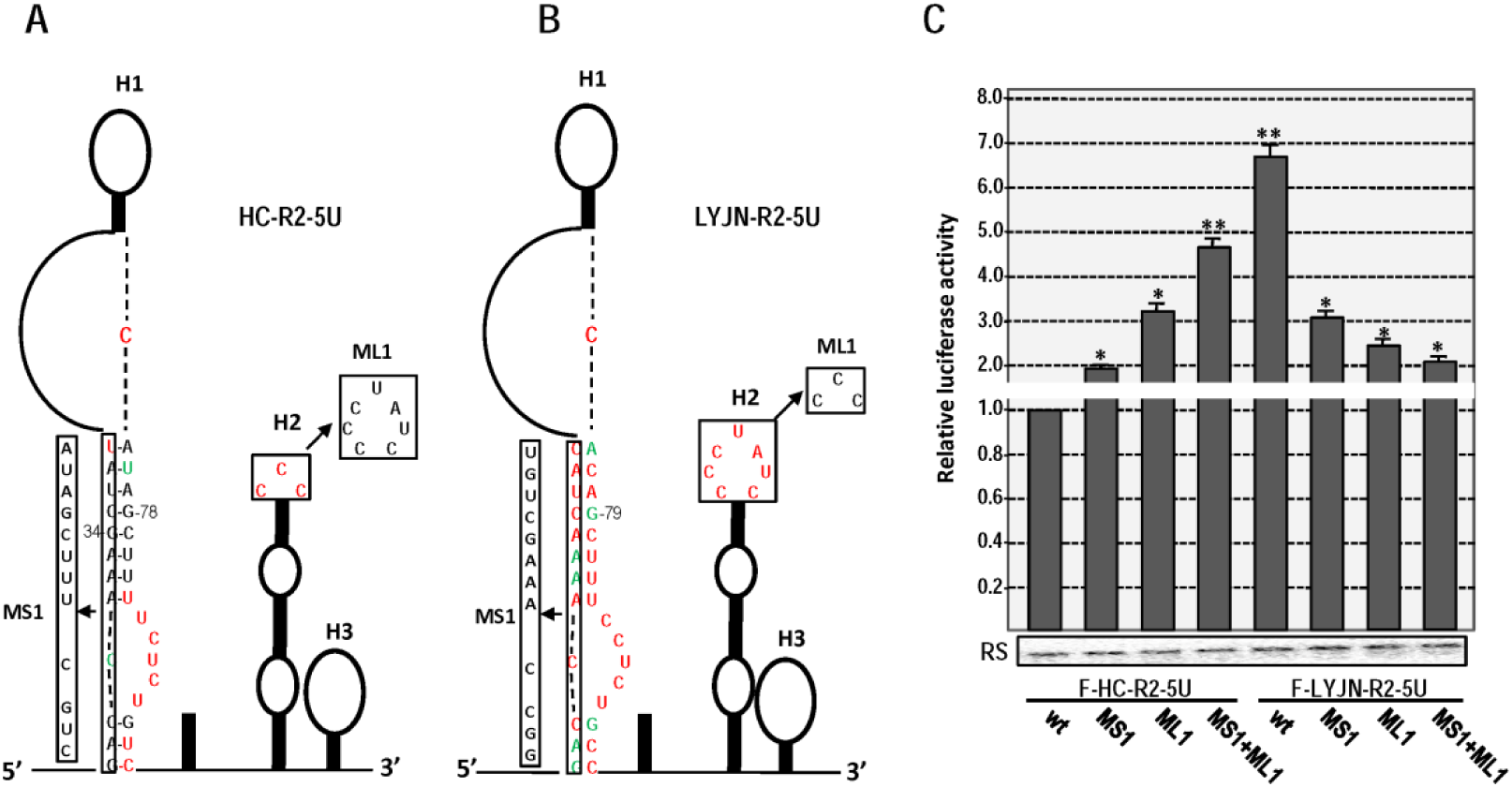
Effect of middle stem of H1 and top loop of H2 in the RNA2 5′-UTRs of WYMV HC and LYJN isolates on translation. A. **Mutations on middle stem of H1 and top loop of H2 in the RNA2 5′-UTRs of WYMV HC isolate.** B. **Mutations on middle stem of H1 and top loop of H2 in the RNA2 5′-UTRs of WYMV LYJN isolate.** C. **Effects of mutations of middle stem of H1 and top loop of H2 in the RNA2 5′-UTRs of WYMV HC and LYJN isolates on translation.** * indicates P < 0.05, ** indicates P < 0.01; RS indicates RNA stability assay after 1.5 h *in vitro* translation.

Except to two regions in H1 and H2 resulting into the variable IRES activity of RNA2 5′-UTR, core regions ensuring the basic IRES activity of RNA2 5′-UTR was also mapped. Top loop and top stem in H1 were mutated to test the effect on IRES activity. For RNA2 5′-UTR of HC isolate, mutant of weaken base pair of top stem in H1 (MS2) had 38% IRES activity of that of wt HC (F-HC-R2-5U), while mutant of nucleotide replacement in top loop of H1 (ML2) only had 20% IRES activity of wt HC (F-HC-R2-5U), which was close to translation level of the control vector (Fig.6A and 6C). For RNA2 5′-UTR of LYJN isolate, mutant of weaken base pair of top stem in H1 (MS2) had 28% IRES activity of that of wt LYJN, while mutant of nucleotide replacement in top loop of H1 (ML2) had 20% IRES activity of that of wt LYJN, which was close to translation level of wt HC (Fig.6B and 6C). Mutation on top loop or top stem in H1 caused similar extent of reduction on IRES activity for both HC and LYJN isolates. It is suggested that nucleotide specificity of top loop and stability of top stem in H1 are essential factor ensuring the basic IRES activity of RNA2 5′-UTR. Firm base pairing status in top stem of H1 can ensure the specific recruitment of corresponding translation initiation factors by specific nucleotide of top loop in H1. Mutant on top loop of H1 in HC isolate almost vanished IRES activity, which implied that top loop of H1was the exclusive site to recruit translation machinery in RNA2 5′-UTR of HC isolate. However, mutant on top loop of H1 in LYJN isolate still had IRES activity as that of RNA2 5′-UTR of HC isolate, which suggested that the top loop of H1 in RNA2 5′-UTR of LYJN was not exclusive site to recruit translation machinery and the top loop of H2 may be the additional site to recruit translation machinery.

**Figure 6.**
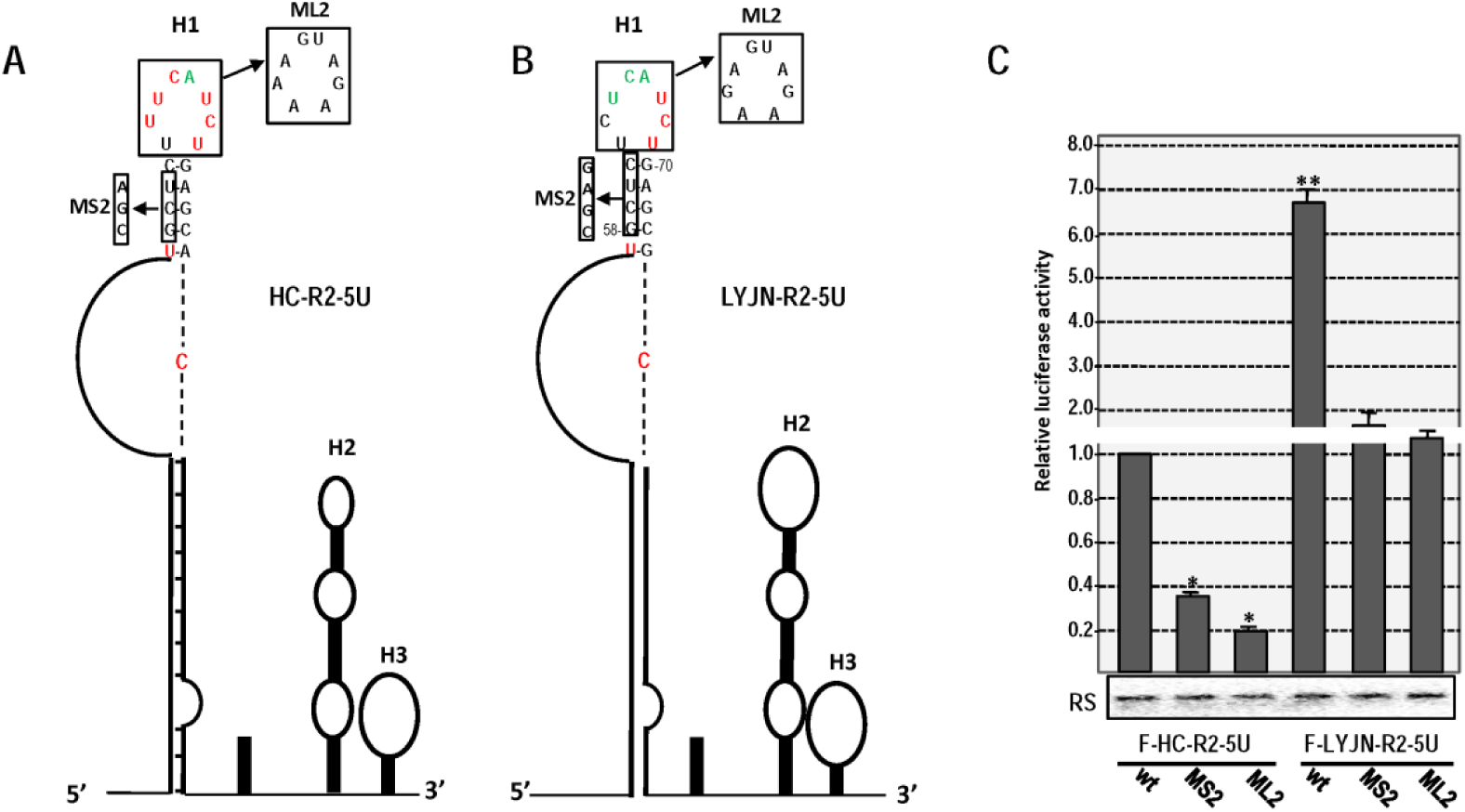
Effect of top loop and top stem of H1 in the RNA2 5′-UTRs of WYMV HC and LYJN isolates on translation. **A. Mutations of top loop and top stem of H1 in the RNA2 5′-UTRs of WYMV HC isolate.** **B. Mutations of top loop and top stem of H1 in the RNA2 5′-UTRs of WYMV LYJN isolate.** **C. Effects of mutations of top loop and top stem of H1 in the RNA2 5′-UTRs of WYMV HC and LYJN isolates on translation.** * indicates P < 0.05, ** indicates P < 0.01; RS indicates RNA stability assay after 1.5 h *in vitro* translation.

### Different RNA2 5′UTR of WYMV had different numbers of recruitment site on wheat eIF4E and the synergistic effect of these recruitment sites was mediated by flexibility of middle stem of hairpin H1

Based on above data, 5′-UTR of WYMV RNA2 had IRES activity mediating cap-independent translation, which is involved in the recruitment on specific translation initiation factor. To identify the specific interaction between RNA2 5′-UTR and translation initiation factor, EMSA was performed (Fig.7). Based on the EMSA signal, RNA2 5′-UTR of HC and LYJN isolates can specifically interact with wheat eIF4E (Wt-eIF4E) but not with peanut eIF4E (Pt-eIF4E) (Fig.7A and 7B). In addition, *trans*-competition assay was performed to test the function of eIF4E on cap-independent translation mediated by RNA2 5′-UTR of WYMV. In the presence of the 1.6 mM dissociative cap analog/Mg^2+^, the IRES activity of the RNA2 5′-UTR of HC was decreased to 5% of that of wt (Fig.7C). This confined IRES activity by the cap analog/Mg^2+^ was rescued to 60% of that of wt through the addition of wheat eIF4E but not peanut eIF4E (Fig.7C). For RNA2 5′-UTR of LYJN, competition of cap analog/Mg^2+^ and remedy of wheat eIF4E on IRES activity had similar pattern (Fig.7D). However, RNA2 5′-UTR of LYJN had higher capacity against competition of cap analog/Mg^2+^ than RNA2 5′-UTR of HC. In the presence of the 1.6 mM dissociative cap analog/Mg^2+^, the IRES activity of the RNA2 5′-UTR of LYJN was decreased to 29% of that of wt (Fig.7D). It is suggested that RNA2 5′-UTR played IRES role through specifically recruiting wheat eIF4E and different RNA2 5′-UTR had different recruitment activity on eIF4E.

**Figure 7.**
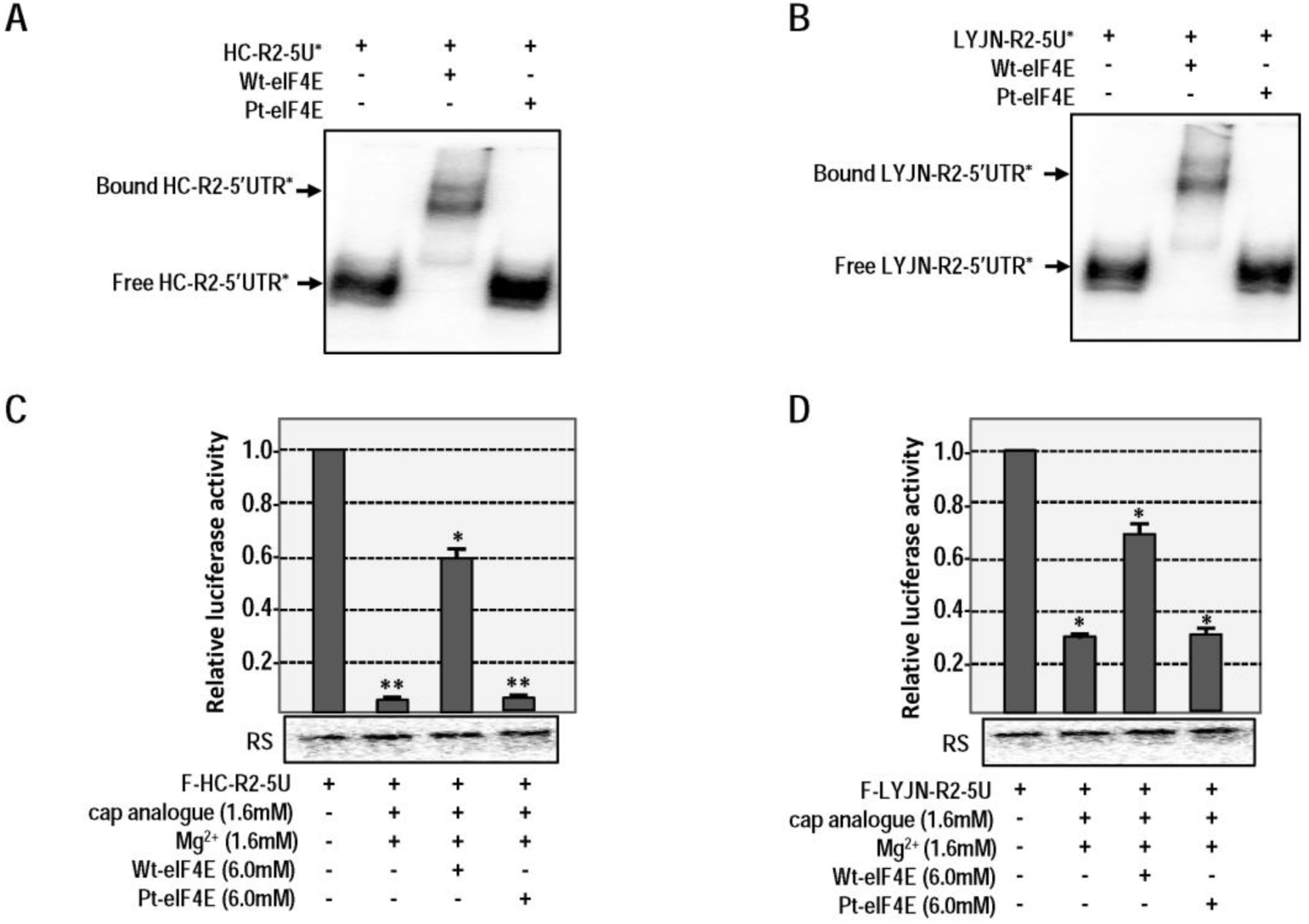
Interaction between eIF4E and RNA2 5′-UTR of WYMV HC or LYJN isolates and its effect on translation. A. **EMSA assay between different eIF4E and RNA2 5′-UTR of WYMV HC isolate.** Wt eIF4E, eIF4E of wheat; Pt eIF4E, eIF4E of peanut. 5′ isotope-labelled RNA2 5′-UTR was indicated by *. B. **EMSA assay between different eIF4E and RNA2 5′-UTR of WYMV LYJN isolate.** C. ***Trans*-competition assay on translation mediated by the RNA2 5′-UTR of WYMV HC isolate from dissociative cap analog and different eIF4E.** * indicates P<0.05, ** indicates P<0.01; RS indicates RNA stability assay after 1.5 h *in vitro* translation. D. ***Trans*-competition assay on translation mediated by the RNA2 5′-UTR of WYMV LYJN isolate from dissociative cap analog and different eIF4E.**

Based on mutagenesis and in vitro translation, it is suggested that RNA2 5′-UTR of HC had one exclusive regions in H1 to recruit translation machinery, while RNA2 5′-UTR of LYJN may have two regions in H1 and H2 to recruit translation machinery (Fig.5 and Fig.6). EMSA and *trans*-competition assay were preformed to test the interaction between these regions and eIF4E. For HC isolate, RNA2 5′-UTR can interact with wheat eIF4E (Fig.7A and Fig.8A). When excess non-isotope labelling HC-R2-5U was added into EMSA assay, the shift signal of isotope labelling wt RNA2 5′-UTR of HC isolate (HC-R2-5U) by the wheat eIF4E disappeared (Fig.8A). However, excess non-isotope labelling mutant of H1 top-loop (HC-R2-5U-ML2) was added into EMSA assay, the shift signal of isotope labelling HC-R2-5U appeared was not affected. *Trans*-competition *in vitro* translation assay also presented the correlative pattern (Fig.8C). It is suggested that top loop of H1 in RNA2 5′-UTR of HC isolate was the exclusive site to specifically recruit wheat eIF4E. For LYJN isolate, RNA2 5′-UTR may have two sites interacting with wheat eIF4E (Fig.7B and Fig.8B). When excess non-isotope labelling LYJN-R2-5U was added into EMSA assay, the shift signal of isotope labelling wt RNA2 5′-UTR of LYJN isolate (LYJN-R2-5U) by the wheat eIF4E disappeared (Fig.8B). When excess non-isotope labelling H1 top-loop mutant (LYJN-R2-5U-ML2) or H2 top-loop mutant (LYJN-R2-5U-ML1) was added into EMSA assay, the shift signal of isotope labelling LYJN-R2-5U was partially remained (Fig.8B). When excess non-isotope labelling dual-mutant of top-loop of H1 and H2 (LYJN-R2-5U-ML1+ML2) was added into EMSA assay, the shift signal of isotope labelling LYJN-R2-5U by the wheat eIF4E was not affected (Fig.8B). *Trans*-competition *in vitro* translation assay also presented the correlative pattern (Fig.8D). It is suggested that top loop of H1 and H2 in RNA2 5′-UTR of LYJN isolate were two sites to specifically recruit wheat eIF4E. *In vitro* structural probing was also performed to map and confirm the interaction between these sites and wheat eIF4E(Fig.9). RNA structure change of RNA2 5′-UTR was analyzed in the absence or presence of wheat eIF4E. In the presence of eIF4E, in-line cleavage of 3′-part of 5′-UTR behind position 70 presented slight weaker (Fig.9A and 9C). Under the integrative consideration of adjacent cleavage pattern and absence/presence of wheat eIF4E, regions with relative weaker in-line cleavage pattern indicated the potential interaction sites with eIF4E. For RNA2 5′-UTR of HC isolate, regions with weaken cleavage pattern in the presence of eIF4E was mainly located at top loop of H1 (Fig.9A and 9B). For RNA2 5′-UTR of LYJN isolate, regions with weaken cleavage pattern in the presence of eIF4E was mainly located at top loop of H1 and H2 (Fig.9C and 9D). It is confirmed that top loop of H1 is the conserved site to recruit wheat eIF4E for RNA2 5′-UTR of HC and LYJN isolate, and top loop of H2 is an additional site for RNA2 5′-UTR of LYJN isolate. *Trans*-competition assay by oligo-nucleotides was performed to test the importance and the potential of these recruitment sites on eIF4E as antiviral target. P1(5-AGATGARA-3) and P2 (5-GATAGGGT-3) was respectively designed to target the top loop of H1 and H2 through complementary base pairing. For RNA2 5′-UTR of HC isolate, only P1 can decrease the IRES activity (Fig.10A). For RNA2 5′-UTR of LYJN, both P1 and P2 can decrease the IRES activity (Fig.10B). It is suggested that oligo-nucleotides blocking recruitment sites on eIF4E could be the effective antiviral agent.

**Figure 8.**
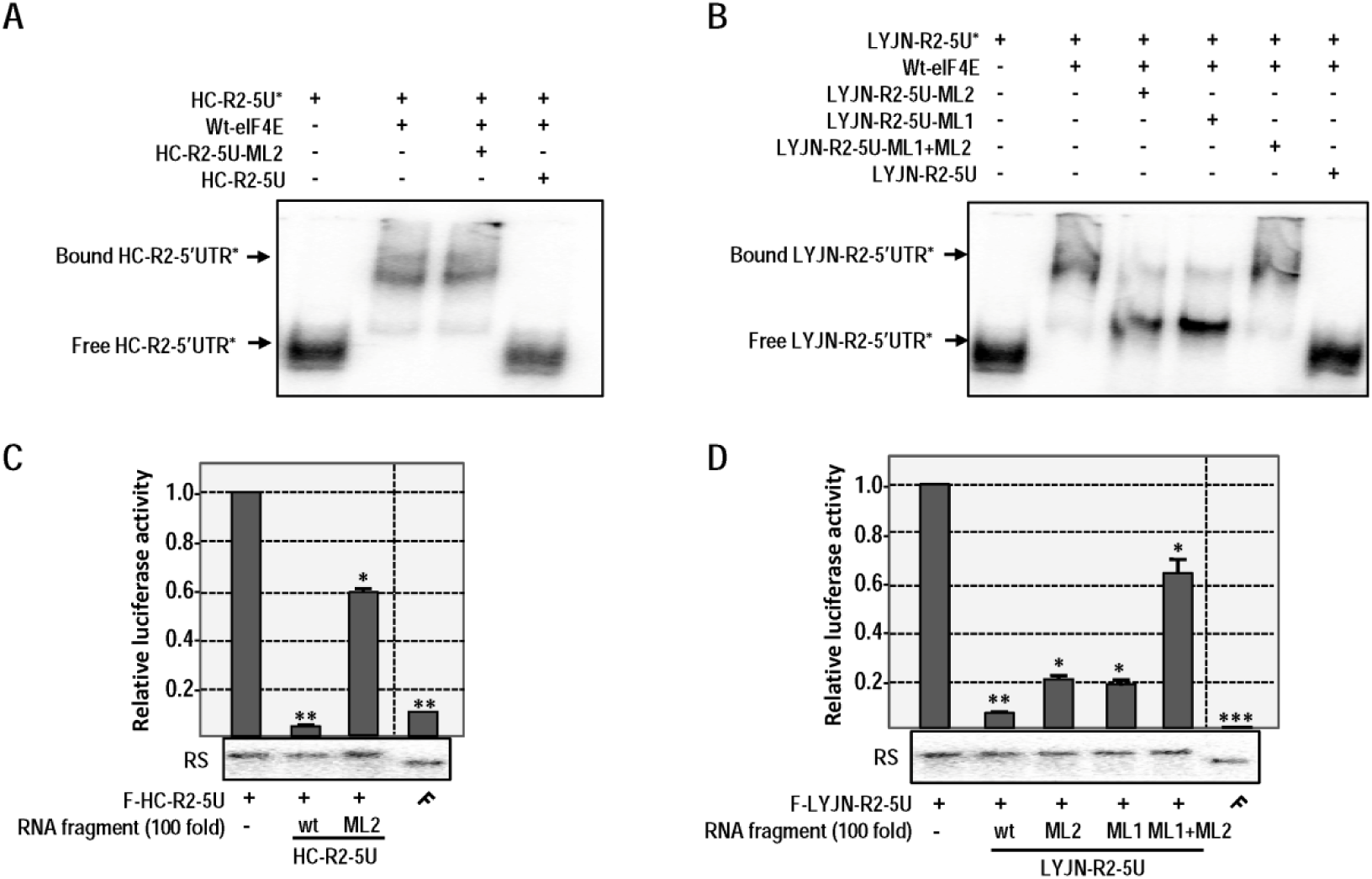
Mapping of interacting regions of RNA2 5′-UTR of WYMV HC and LYJN isolates with wheat eIF4E. **A. EMSA assay among isotope labelled RNA2 5′-UTR of WYMV HC isolate, wheat eIF4E and different RNA fragments derived from RNA2 5′-UTR of HC isolates.** **B. EMSA assay among isotope labelled RNA2 5′-UTR of WYMV LYJN isolate, wheat eIF4E and different RNA fragments derived from RNA2 5′-UTR of LYJN isolates.** **C. *Trans*-competition assay on translation mediated by the RNA2 5′-UTR of WYMV HC isolate from wt RNA2 5′-UTR or its mutants of HC isolate.** * indicates P < 0.05, ** indicates P < 0.01, *** indicates P < 0.001; RS indicates RNA stability assay after 1.5 h *in vitro* translation. **D. *Trans*-competition assay on translation mediated by the RNA2 5′-UTR of WYMV LYJN isolate from wt RNA2 5′-UTR or its mutants of LYJN isolate.**

**Figure 9.**
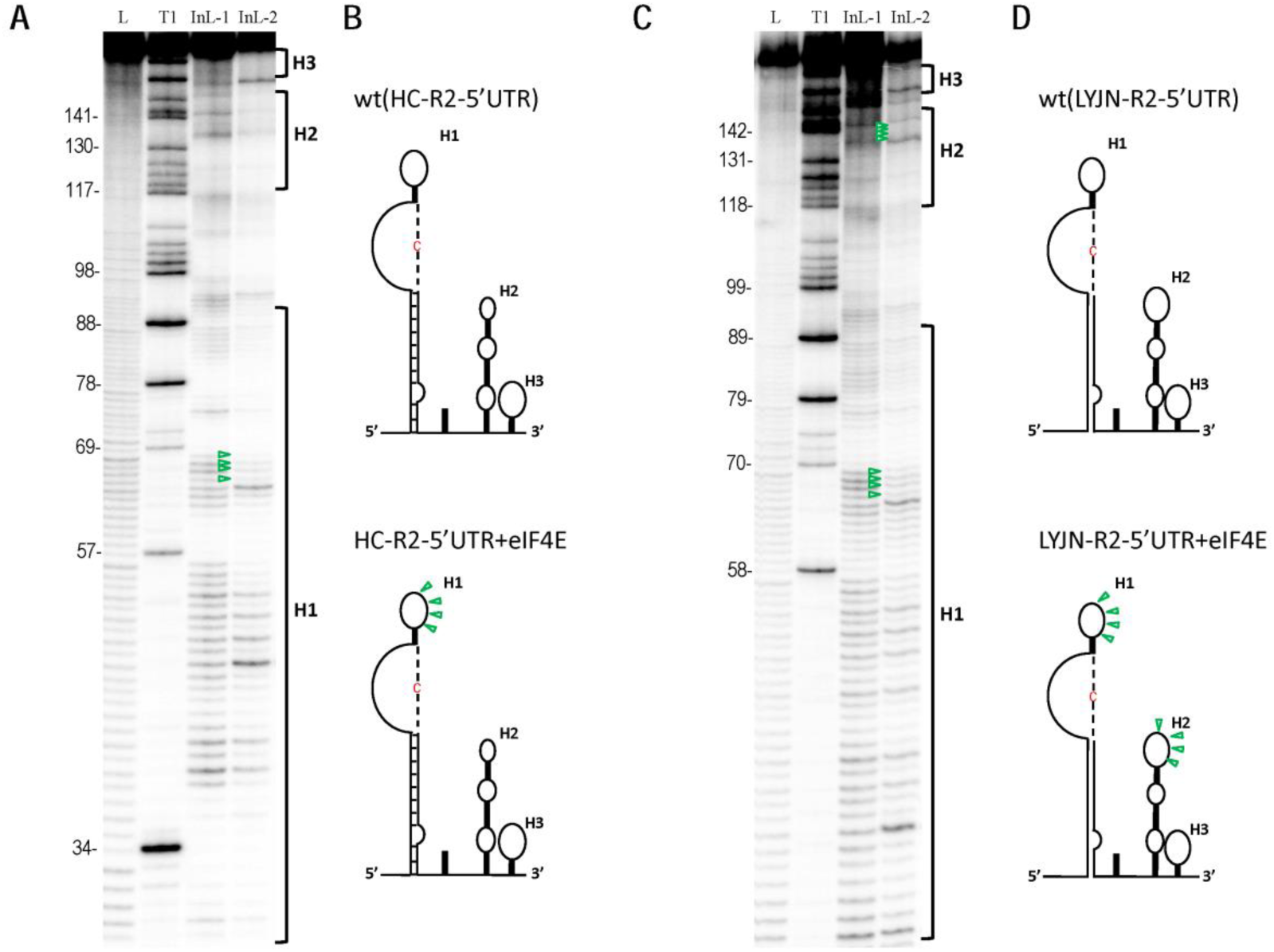
Structure probing of the RNA2 5′-UTR of WYMV HC and LYJN isolates in the presence of the wheat eIF4E. A. **In-line probing of the RNA2 5′-UTR of WYMV HC isolate in the presence of the wheat eIF4E.** The legend is similar to that of Figure 4A. Green hollow triangles point to the weaker cleavage sites within the RNA2 5′-UTR in the presence of the wheat eIF4E versus the absence of the wheat eIF4E. InL-1, In-line cleavge reaction of RNA2 5′-UTR of WYMVHC isolate; InL-2: In-line cleavge reaction of RNA2 5′-UTR of WYMVHC isolate in the presence of the wheat eIF4E. B. **RNA structure of RNA2 5**′**-UTR of WYMVHC isolate in the absence or the presence of the wheat eIF4E.** Green hollow triangles point to the weaker cleavage sites within the RNA2 5′-UTR in the presence of the wheat eIF4E. C. **In-line probing of the RNA2 5′-UTR of WYMV LYJN isolate in the presence of the wheat eIF4E.** InL-1, In-line cleavge reaction of RNA2 5′-UTR of WYMV LYJN isolate; InL-2: In-line cleavge reaction of RNA2 5′-UTR of WYMV LYJN isolate in the presence of the wheat eIF4E. D. **RNA structure of RNA2 5′-UTR of WYMV LYJN isolate in the absence or the presence of the wheat eIF4E.**

**Figure 10.**
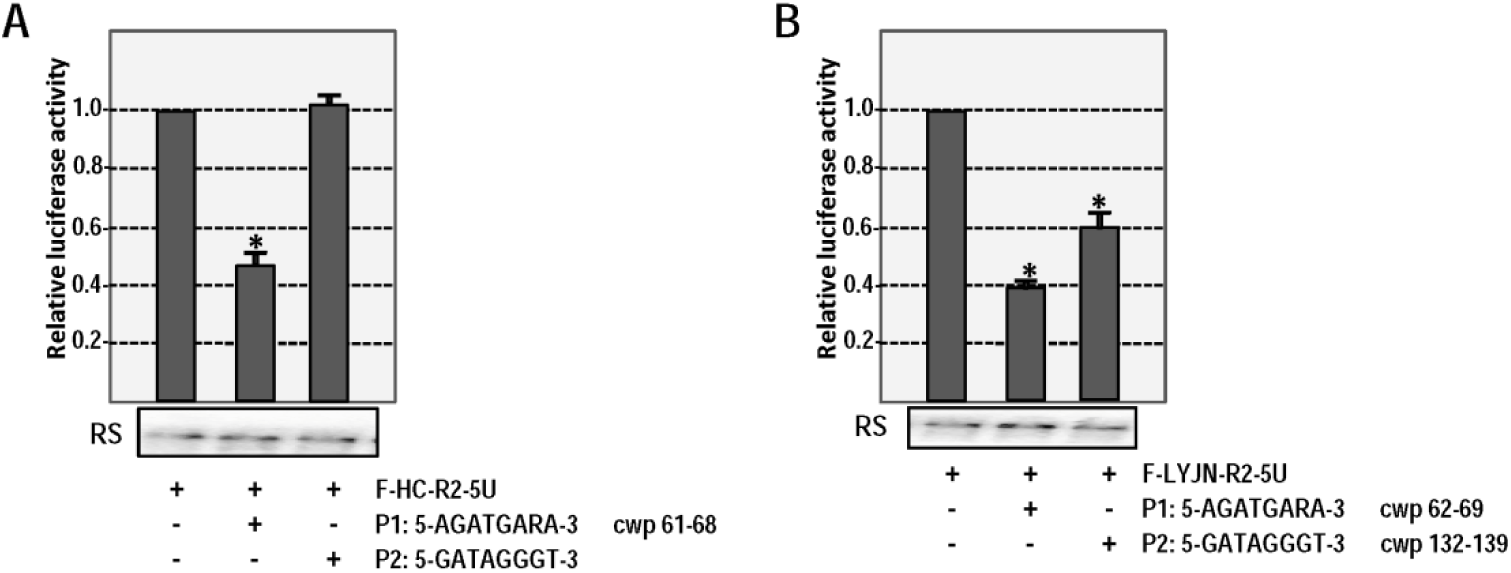
*Trans*-competition from oligonucleotides on translation mediated by RNA2 5′-UTR of WYMV HC and LYJN isolates. A. ***Trans*-competition from different 8 nt oligonucleotides on translation mediated by the RNA2 5′-UTR of WYMV HC isolate**. cwp, complementary with the position. **B. *Trans*-competition from different 8 nt oligonucleotides on translation mediated by the RNA2 5′-UTR of WYMV LYJN isolate.**

Taken together, different 5′-UTR of WYMV RNA2 presented remarkable variable IRES activity, which displayed marked contrast to the conserved IRES activity of 5′-UTR of WYMV RNA1. Variable IRES activity of WYMV RNA2 was mainly determined by numbers of recruitment site on wheat eIF4E. Top loop of H1 was the conserved site interacting with wheat eIF4E in both HC and LYJN isolate (Fig.11A), while top loop of H2 was an additional site interacting with wheat eIF4E in LYJN isolate (Fig.11B). Furthermore, top loop of H1 and H2 in RNA2 5′-UTR of LYJN can synergistically recruit eIF4E to greatly enhance IRES activity, because loosened middle stem of H1 can allow top loop of H1 and H2 to be adjacent to each other (Fig.11B).

**Figure 11.**
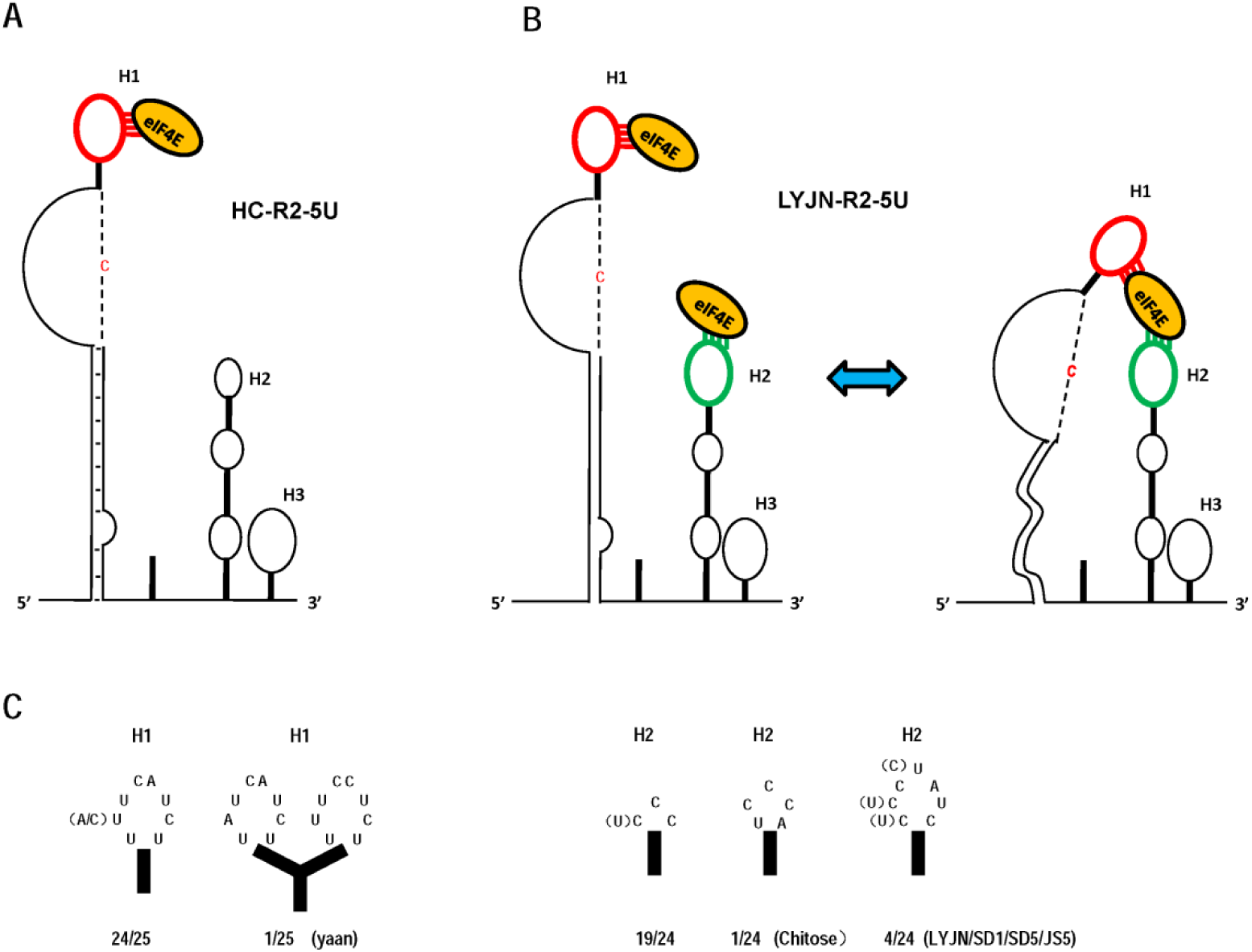
Characteristics of core *cis*-elements of IRES within 5′-UTR of WYMV RNA2 and specific interaction with the wheat eIF4E. **A. Interaction between eIF4E and IRES of 5′-UTR of RNA2 of WYMV HC isolate.** H1, hairpin1; H2, hairpin 2; H3, hairpin 3; Red lines indicate the interaction of H1 (red loop) and eIF4E. **B. Dynamic interaction between eIF4E and IRES of 5′-UTR of RNA2 of WYMV LYJN isolate.** Red lines indicate the interaction of H1 (red loop) and eIF4E; Green lines indicate the interaction of H2 (green loop) and eIF4E. **C. Characteristic of top loop of H1 and H2 among RNA2 5′-UTR of different WYMV isolates.** yaan: RNA2 of WYMV yaan isolate (AJ2424490); Chitose: RNA2 of WYMV Chitose isolate (AB627822). SD1: RNA2 of WYMV SD1 isolate (KY354399); SD5: RNA2 of WYMV SD5 isolate (KY354406); JS5: RNA2 of WYMV JS5 isolate (KY354403).

RNA2 of 24 isolates of reported 25 isolates had conserved top loop of H1. RNA2 of 19 isolates had tri-nucleotide top loop of H2 as HC isolate, and 4 isolates had hepta-nucleotide top loop of H2 as LYJN isolate (Fig.11C). It is suggested that top loop of H1 in WYMV RNA2 was conserved and top loop of H2 in WYMV RNA2 mainly existed as two types. RNA2 of Chitose isolate had penta-nucleotide top loop of H2, whose IRES activity maybe intermediate state between HC and LYJN isolate (Fig.11C). RNA2 of yaan isolate had two similar top loops of H1 without containing top loop of H2 (Fig.11C), which may represent another pattern of high IRES activity due to the synergistic effect of two similar top loop of H1 to recruit eIF4E.

## DISCUSSION

### IRESes variation among different species, viruses, isolates of same virus or genomic RNAs within one virus

Up to date, IRESes have been identified in viruses, yeast, fruit fly and human cellular mRNA (Gilbert *et al,* 2007; Kieft, 2008; Xia & Holcik, 2009; Weingarten-Gabbay *et al,* 2016). All IRESes can mediate cap-independent translation, but they have different sequences, structures and mechanisms involved in the requirement of translation initiation factors even and/or IRES trans-acting factors (ITAFs) (Sweeney *et al,* 2012; Lozano & Mart′ınez-Salas, 2015; zhang *et al,* 2015). IRESes of animal viruses had more complicated structure than that of other resources such as plant viruses and eukaryotic cellular mRNA, of which IRESes had weaker structure with few hairpins even no obvious structure. It is suggested that IRESes of animal viruses may undergo different evolution pathway from that of other resource. However, structure complexity of IRESes was not correlated with corresponding high activity. IRESes with weaker structure still had high IRES activity (Xia & Holcik, 2009), which was also shown in this study. Except the structure difference, IRESes of different viruses had different mechanism to recruit translation machinery mainly involved in requirement on different translation initiation factors for the final recruitment of 40S ribosomal subunit.

In addition to the different IRESes among different viruses, IRESes of different isolates of same virus also had different structure characteristic and variable activity as shown in this study. 5′-UTR of different WYMV RNA2 had remarkable difference of IRES activity correlated with the different structure characteristic (Fig.11). Furthermore, IRESes in different genomic RNAs within one virus also had different evolution pattern. IRESes in WYMV RNA1 and RNA2 had different structure characteristic and evolution pattern. IRESes in WYMV RNA1 were conserved among different isolates (Geng *et al,* 2020), while IRESes in WYMV RNA2 were variable among different isolates in this study. Taken together, IRES variation among different species, viruses, isolates of same virus or genomic RNAs within one virus suggested that evolution of IRESes may be related with origin resource, RNA type, circumstance to play role even the characteristic of proteins expressed via IRES.

### Perception of virus evolution from the study of WYMV IRESes

As obligate parasite, the basic principle of virus evolution should be to adopt the host in different condition and keep the balance between self-duplication and harmfulness on host. WYMV had two genomic RNAs, whose IRES presented different evolution pattern. IRESes of different WYMV RNA1 had conserved dynamic equilibrium state of RNA tertiary structure, regulating the optimal not maximum translation level of viral protein encoded by RNA1 (Geng *et al,* 2020). However, IRESes of different WYMV RNA2 had well marked different activity and structural characteristic in this study. This different evolution pattern may be related with the characteristic of viral proteins encoded by WYMV RNA1 or RNA2. WYMV RNA1 encoded a polyprotein, from which Nib with RNA-dependent RNA polymerase activity was produced and responsible for replication of progeny viral RNAs (Namba *et al,* 1998; Geng *et al,* 2019). dsRNA stage during replication of RNA viruses can trigger RNA-induced silencing complex (RISC) of host to cleavag themselves (Zamore *et al,* 2000). If dsRNA are produced too much or too fast, RISC will be easily triggered to cleavage dsRNA, which will restrict self-duplication of virus. So optimal replication of virus is to keep the balance of self-duplication and low harmfulness on host, which may not trigger the cleavage of RISC. In this case of WYMV RNA1, the evolution pattern of dynamic equilibrium state IRESes was matched with the requirement of the optimal not maximum level of NIb (Geng *et al,* 2020). WYMV RNA2 encoded two proteins responsible for WYMV transmission via fungus, formation of replication locus and possible gene-silencing suppressor (Dessens & Meyer, 1996; Tatineni *et al,* 2012; Sun *et al,* 2014; Xie *et al,* 2019). So high level of proteins encoded by WYMV RNA2 is helpful for WYMV transmission and self-duplication within replication locus. The evolution tendency of IRES in WYMV RNA2 was matched with the requirement of high level expression of viral proteins encoded by RNA2. Based on characteristic about conservative IRES activity in WYMV RNA1 and variable IRES activity in WYMV RNA2, it is implied an equilibrium status of “offensive and defensive tactics” during evolution and survive of RNA viruses. In addition, IRES of WYMV RNA2 specifically recruited wheat eIF4E, explaining from one side why wheat was the only natural host of WYMV. It provided new evidence for eIF4E is one of core factors of non-host resistance (Nieto *et al,* 2011; Baruah *et al,* 2020). Gene editing on wheat eIF4E may be practical strategy to breed resistant wheat against WYMV (Bastet *et al,* 2017).

## Material and methods

### Construction of plasmids and preparation of DNA fragments

All plasmids were constructed based on the firefly luciferase reporter construct pT7-F-3′-UTRssp vector (Wang *et al,* 2017) via PCR amplification, enzyme digestion and ligation. All plasmids were confirmed by DNA sequencing. Detailed information about plasmid construction and corresponding in vitro transcripts are shown in Supplementary Table1.

DNA fragments were amplified via PCR to be the template for making corresponding *in vitro* transcripts, which was used for EMSA, In-line probing and *trans*- competition assay. Detailed information was shown in Supplementary Table2.

### *In vitro* translation

In vitro translation assays were performed as previously described (Wang *et al,* 2017). Briefly, the *in vitro*-synthesized RNA transcripts (3 pmol) from designated translation reporter constructs were used for a 25 µL translation reaction volume using wheat germ extracts (WGE; Promega) according to the manufacturer’s instructions. The luciferase activity was measured by using a luciferase assay reporter system (Promega) and a Modulus microplate multimode reader (Turner BioSystems). At least three independent in vitro translation assays were performed for each construct. Standard errors were calculated in Microsoft Excel. Statistical analyses of the significance of in vitro translation assays were performed using SPSS or t-test, as indicated (George & Mallery, 2016).

### RNA stability assay

RNA stability assays were performed after 1.5 h of in vitro translation using northern blotting. Sequences of specific probe for northern blotting are 5-acgtgatgttcacctcgatatgtgcatctg-3, which is complemented with the coding region of the firefly luciferase (FLuc).

### eIF4E expression and purification

The coding sequences of wheat eIF4E (KX467331) or peanut eIF4E (HE985069) was amplified by RT-PCR and inserted into pEHis-TEV vector through enzyme site *Bam*H I and *Sal* I. The positive plasmid was transformed into *E.coli* Rosetta. And eIF4E proteins were expressed under the induction of 0.5 M IPTG at 37 ℃ for 5 h. Samples were separated by SDS-PAGE. The gel band corresponding to the 24 kDa of eIF4E protein were cut and putted into dialysis bag. After 2 h electrophoresis in SDS-PAGE buffer, the gel in dialysis bag was removed and dialysis bag were dialyzed for 24 h followed by split charging of the eIF4E proteins. The purified wheat eIF4E and peanut eIF4E were used in following In-line structure probing, electrophoretic mobility shift assay (EMSA), and trans-competition assay.

### In-line structure probing

In-line structure probing was performed as previously described (Yuan *et al,* 2012). Briefly, RNA fragments of the WYMV-HC RNA2 5′-UTR (positions 1-171 nt) or the WYMV-LYJN RNA2 5′-UTR (positions 1-172 nt) were separately 5′-end-labelled with [γ-^32^P] ATP and denatured at 75 ℃ followed by slowly cooling to 25 ℃. RNA was incubated at 25 ℃ in 50 mM Tris-HCl [pH 8.5] and 20 mM MgCl_2_ for 14 h. To identify potential RNA-protein interaction, unlabeled candidate eIF4E were co-incubated with isotope-labelled RNA fragments. Samples were separated by 8% polyacrylamide gel electrophoresis (8M urea) alongside a hydroxide-generated RNA cleavage ladder and RNase T1 digestion product on labelled RNA fragments. Then, the gels were dried and exposed to a phosphorimager screen, followed by detection with the Typhoon FLA-7000 (GE Healthcare). At least two independent in-line probing assays were performed for each fragment. Signals of the in-line probing reaction were analyzed as previously described (Geng *et al,* 2020).

### Electrophoretic mobility shift assay (EMSA)

EMSA was used to test the interaction between eIF4E and 5′-UTR of WYMV RNA2 or its mutants in this study. 5′-end-labeled 5′-UTR of WYMV RNA2 (AF041041 or KX258949) or unlabeled other RNA fragments for competition assay was put into 75 ℃ with natural temperature reduction for denaturation-natural refolding treatment until reaction temperature reaches room temperature (25 °C). Subsequently, denaturation-natural refolding treatment 5′-end-labeled 5′-UTR of WYMV RNA2 and 50 fold of eIF4E without or with 500 fold of unlabeled other RNA fragments for competition assay was added into the final volumn of 10.0 μL, and 10.0 μL2×RNA binding buffer (10 mM HEPES pH7.9, 200 mM KCl, 10 mM MgCl_2_, 7.6% glycerol) was added followed by reaction at 25°C for 0.5 h, and 5.0 μL RNA-protein EMSA loading buffer (2×RNA binding buffer 2.5 μL, 100% glycerol 2.5 μL, Xylene cyanol FF 0.001g and Bromophenol blue 0.001g) were added to the reaction solution. Reaction products were separated by 5% native polyacrylamide gel electrophoresis. Then, the gels were dried and exposed to a phosphorimager screen, followed by detection with the Typhoon FLA-7000 (GE Healthcare).

### *Trans*-competition assay

The *trans*-competition assay was performed based on the *in vitro* translation assay to test the effect of the additional competing items including 1.6 mM cap analogue/Mg^2+^, 6 mM eIF4E, 100-foldof RNA fragments or 8 μM oligonucleotides.

## Data availability

The data that support the findings of this study are available from the corresponding author upon reasonable request.

## Acknowledgments

We are grateful to Prof. Chenggui Han (China Agricultural University) for providing the plasmid containing RNA2 of WYMV HC isolate. The work was funded and supported by National Natural Science Foundation of China (31872638, 32072382, 31670147).

## Author contributions

G.G. contributed to the design of the study, in-line, in vitro translation assay, EMSA and drafting the manuscript. C.Y. contributed to in vitro translation assay and data analysis. X. L. contributed to data analysis. X.Y. and K. S. contributed to the design of the study, data analysis and drafting the manuscript. All authors read and approved the final manuscript.

## Competing interests

The authors declare that they have no competing interests.

## References

1. Baird SD, Turcotte M, Korneluk RG, Holcik M (2006) Searching for ires. RNA 12: 1755–1785.

2. Baruah A, Sivalingam PN, Fatima U, Senthil-Kumar M (2020) Non-host resistance to plant viruses: What do we know? Physiol. Mol. Plant Pathol. 111: 101506.

3. Bastet A, Christophe Robaglia C, Gallois JL (2017) eIF4E resistance: natural variation should guide gene editing. Trends Plant Sci. 22: 411–419.

4. Boerneke MA, Dibrov SM, Gu J, Wyles DL, Hermann T (2014) Functional conservation despite structural divergence in ligand-responsive RNA switches. Proc. Natl. Acad. Sci. U.S.A. 111: 15952–15957.

5. Dessens JT, Meyer M (1996) Identification of structural similarities between putative transmission proteins of Polymyxa and Spongospora transmitted bymoviruses and furoviruses. Virus Genes 12: 95–99.

6. Dibrov SM, Ding K, Brunn ND, Parker MA, Bergdahl BM, Wyles DL, Hermann T (2012) Structure of a Hepatitis C virusRNA domain in complex with a translation inhibitor reveals a binding mode reminiscent of riboswitches. Proc. Natl. Acad. Sci. U.S.A. 109: 5223–5228.

7. Fern′andez, I S, Bai XC, Murshudov G, Scheres SH, Ramakrishnan V (2014) Initiation of translation by cricket paralysis virus IRES requires its translocation in the ribosome. Cell 157: 823–831.

8. Filbin ME, Vollmar BS, Shi D, Gonen T, Kieft JS (2013) HCV IRES manipulates the ribosome to promote the switch from translation initiation to elongation. Nat. Struct. Mol. Biol. 20: 150–158.

9. Gasparian AV, Neznanov N, Jha S, Galkin O, Moran JJ, Gudkov AV, Komar AA (2010) Inhibition of encephalomyocarditis virusand poliovirus replication by quinacrine: implications for the design and discovery of novel antiviral drugs. J. Virol. 84: 9390–9397.

10. Geng GW, Yu CM, Li XD, Yuan XF (2019) Variable 3′polyadenylation of Wheat yellow mosaic virus and its novel effects on translation and replication. Virol. J. 16: 23.

11. Geng GW, Yu CM, Li XD, Yuan, XF (2020) A unique internal ribosome entry site representing a dynamic equilibrium state of RNA tertiary structure in the 5′-UTR of Wheat yellow mosaic virus RNA1. Nucleic Acids Res. 48: 390–404.

12. Geng GW, Yu CM, Wang DY, Gu K, Shi KR, Li XD, Tian YP, Yuan XF (2017) Evolutional analysis of two new isolates of Wheat yellow mosaic virus from Shandong Province, China (in chinese). Acta Phytopathol.Sinica. 47: 348–356.

13. George D, Mallery P (2016) IBM SPSS Statistics 23 Step by Step: A Simple Guide and Reference. Routledge NY and OX.

14. Gilbert WV, Zhou K, Butler TK, Doudna JA (2007) Cap-independent translation is required for starvation-induced differentiation in yeast. Science 317: 1224–1227.

15. Hashem Y, Des Georges A, Dhote V, Langlois R, Liao HY, Grassucci RA, Frank J (2013) Hepatitis-C-virus-like internal ribosome entry sites displace eIF3 to gain access to the 40S subunit. Nature 503: 539–543.

16. Jang SK, Kräusslich HG, Nicklin MJ, Duke GM, Palmenberg AC, Wimmer EA (1988) Segment of the 5′nontranslated region of Encephalomyocarditis virusRNA directs internal entry of ribosomes during in vitro translation. J. Virol. 62: 2636–2643.

17. Kieft JS (2008) Viral IRES RNA structures and ribosome interactions. Trends Biochem. Sci. 33: 274–283.

18. Lozano G, Mart′ınez-Salas E (2015) Structural insights into viral IRES-dependent translation mechanisms. Curr. Opin. Virol. 12: 113–120.

19. Marcotrigiano J, Gingras AC, Sonenberg N, Burley SK (1997) Cocrystal structure of the messenger RNA 5′ cap-binding protein (eIF4E) bound to 7-methyl-GDP. Cell 89: 951–961.

20. Namba S, Kashiwazaki S, Lu X, Tamura M, Tsuchizaki T (1998) Complete nucleotide sequence of Wheat yellow mosaic bymovirus genomic RNAs. Arch.Virol. 143: 631–643.

21. Nicholson BL, White KA (2011) 3′ Cap-independent translation enhancers of positive-strand RNA plant viruses. Curr. Opin. Virol. 1: 373–380.

22. Nieto C, Rodríguez-Moreno L, Rodríguez-Hernández AM, Aranda MA, Truniger V (2011) Nicotiana benthamiana resistance to non-adapted Melon necrotic spot virus results from an incompatible interaction between virus RNA and translation initiation factor 4E. Plant J. 66: 492–501.

23. Pelletier J, Sonenberg N (1988) Internal initiation of translation of eukaryotic mRNA directed by a sequence derived from poliovirus RNA. Nature 334: 320–325.

24. Pestova TV, Lorsch JR, Hellen CUT (2007) The mechanism of translation initiation in eukaryotes. In: Mathews MB, Soneberg N, Hershey JWB, (eds). Translational Control in Biology and Medicine. Cold Spring Harbor Laboratory Press, NY, pp. 87–128.

25. Seth PP, Miyaji A, Jefferson EA, Sannes-Lowery KA, Osgood SA, Propp SS, Ranken R, Massire C, Sampath R, Ecker DJ (2005) SAR by MS: discovery of a new class of RNA-binding small molecules for the Hepatitis C virus: internal ribosome entry site IIA subdomain. J. Med. Chem. 48: 7099–7102.

26. Simon AE, Miller WA (2013) 3′ cap-independent translation enhancers of plant viruses. Annu. Rev. Microbiol. 67: 21–42.

27. Sonenberg N, Hinnebusch AG (2009) Regulation of translation initiation in eukaryotes: mechanisms and biological targets. Cell 136:731–745.

28. Stupina VA, Meskauskas A, Mccormack JE, Yingling YG, Shapiro BA, Dinman JD, Simon AE (2008) The 3′ proximal translational enhancer of Turnip crinkle virus binds to 60S ribosomal subunits. RNA 14: 2379–2393.

29. Stupina VA, Yuan XF, Meskauskas A, Dinman JD, Simon AE (2011) Ribosome binding to a 5′ translational enhancer is altered in the presence of the 3′UTR in cap-independent translation of Turnip Crinkle Virus. J. Virol. 85: 4638–4653.

30. Sun L, Andika IB, Shen J, Yang D, Chen J (2014) The P2 of Wheat yellow mosaic virus rearranges the endoplasmic reticulum and recruits other viral proteins into replication-associated inclusion bodies. Mol. Plant Pathol. 15: 466–478.

31. Sweeney TR, Dhote V, Yu Y, Hellen CU (2012) A distinct class of internal ribosomal entry site in members of the Kobuvirus and proposed Salivirus and Paraturdivirus genera of the Picornaviridae. J. Virol. 86: 1468–1486.

32. Tatineni S, Qu F, Li R, Morris TJ, French R (2012) Triticum mosaic poacevirus enlists P1 rather than HC-Pro to suppress RNA silencing-mediated host defense. Virology 433: 104–115.

33. Ungureanu NH, Cloutier M, Lewis SM, De Silva N, Blais JD, Bell JC, Holcik M (2006) Internal ribosome entry site-mediated translation of apaf-1, but not xiap, is regulated during uv-induced cell death. J. Biol. Chem. 281: 15155–15163.

34. Wang D, Yu C, Liu S, Wang G, Shi K, Li X, Yuan X (2017) Structural alteration of a BYDV-like translation element (BTE) that attenuates p35 expression in three mild Tobacco bushy top virus isolates. Sci. Rep. 7: 4213.

35. Weingarten-Gabbay S, Elias-Kirma S, Nir R, Gritsenko AA, Stern-Ginossar N, Yakhini Z, Weinberger A, Segal E (2016) Systematic discovery of cap-independent translation sequences in human and viral genomes. Science 351: 240–240.

36. Xia X, Holcik M (2009) Strong eukaryotic IRESs have weak secondary structure. PLoS ONE 4: e4136.

37. Xie L, Song X, Liao Z, Wu B, Yang J, Zhang H, Hong J (2019) Endoplasmic reticulum remodeling induced by Wheat yellow mosaic virus infection studied by transmission electron microscopy. Micron 120: 80–90.

38. Xue S, Tian S, Fujii K, Kladwang W, Barna M (2015) Rna regulons in hox 5′UTRs confer ribosome specificity to gene regulation. Nature 517: 33–38.

39. Yuan X, Shi K, Simon AE (2012) A local, interactive network of 3′ RNA elements supports translation and replication of Turnip crinkle virus. J. Virol. 86: 4065–4081.

40. Zamore P, Tuschl T, Sharp P, Bartel D (2000) RNAi: double-stranded RNA directs the ATP-dependent cleavage of mRNA at 21 to 23 nucleotide intervals. Cell 101: 25–33.

41. Zhang J, Roberts R, Rakotondrafara AM (2015) The role of the 5′untranslated regions of Potyviridae in translation. Virus Res. 206: 74–81.

